# A modular and flexible open source cell incubator system for mobile and stationary use

**DOI:** 10.1101/2024.03.27.587077

**Authors:** Jens Duru, Benedikt Maurer, Tobias Ruff, Julian Hengsteler, Sophie Girardin, János Vörös, Stephan J. Ihle

## Abstract

Culturing living cells *in vitro* requires the maintenance of physiological conditions for extended periods of time. Here, we introduce a versatile and affordable incubation system, addressing the limitations of traditional incubation systems. Conventionally, stationary cell incubators maintain constant temperature and gas levels for *in vitro* cell culturing. Combining such incubators with additional lab equipment proves challenging. The presented platform offers modularity and adaptability, enabling customization to diverse experimental needs. The system includes a main unit with a user-friendly interface as well as an interchangeable incubation chamber. We present two incubation chambers targeting two completely different use cases. The first chamber, named “inkugo” facilitates the transportation of cells for up to two hours without external power and for more than a day without an external CO_2_ source. The second chamber termed “inkubox” was designed to enable continuous electrophysiological recordings. Recordings from up to four neural cultures growing on high-density microelectrode arrays can be performed in parallel. The system’s unique feature lies in its separability of control and incubation components, allowing one control unit to manage various custom chambers. The design’s simplicity and the use of widely accessible components make the here proposed incubation system replicable for any laboratory. This platform fosters collaboration and experimentation in both decentralized and traditional laboratory settings, making it an invaluable addition to any cell culturing pipeline.

**Specifications table**

|  |  |
| --- | --- |
| **Hardware name** | inkugo and inkubox |
| **Subject area** | Biological sciences |
| **Hardware type** | Biological sample handling and preparation |
| **Closest commercial analog** | cell incubators for stationary and/or mobile use |
| **Open source license** | CC BY-SA 4.0 |
| **Cost of hardware** | approx. 750 USD |
| **Source file repository** | https://downloads.lbb.ethz.ch/2024_Duru_et_al_INKUBOX/ |

## 1 Hardware in context

Sustaining living cells in an *in vitro* cell culturing environment mandates the perpetual maintenance of physiological conditions in said environment. Conventionally, the requirements for prolonged *in vitro* cultivation are fulfilled by stationary cell incubators. These systems usually uphold a temperature of 37 °C. In addition, a CO_2_ concentration of 5 % is required [1]. Incubators are considered a standard apparatus within any cell laboratory. Nevertheless, difficulties emerge when additional conditions are required for such incubators, for example, the need for portability or the interest in having additional laboratory equipment interact with the cell cultures without negatively affecting the culture’s environment. The deficiency of adaptability and flexibility in commercially available stationary incubators poses numerous obstacles in experimental designs.

If external laboratory equipment, such as an electrophysiological system, cannot easily be combined with a commercial incubator, time limitations can arise. Placing a living culture within a recording unit outside an incubator drastically limits the available time for experiments, since a drop in viability can be expected when *in vitro* cultures are exposed to non-physiological conditions. Moreover, an experimental design where cultures are moved continuously back and forth between an incubator and a recording system exposes the cell culture to repeated thermal as well as mechanical shock. Such shocks can have an adverse effect on cell behavior including culture viability [2, 3]. Another challenge arises with the need to relocate live cell cultures from one facility to another without impairing their viability.

The advent of cheap and widely available processing units such as Arduinos [4] and Raspberry Pis [5] have accelerated the development of research tools. Such open-source units are used to control a plethora of laboratory devices ranging from biosensing [6] to microscopy [7]. In addition, the progress made in the field of table-top 3D printing has opened new avenues for scientists to produce such tools [8, 9]. Such 3D printers can be used to print necessary parts required for laboratory equipment and the replication of new tools.

The field of bottom-up neuroscience[10] has profited from the ever-increasing quality of new 3D printing and microfluidic tools. By now such devices can be used to control axonal growth with high fidelity in neuronal cultures [11, 12] even with highly sensitive cells such as human-induced pluripotent stem cell (hiPSC)-derived neurons [13, 14]. While these approaches introduce a high level of control to the cultures themselves, such control is only useful, if interaction with such cell cultures occurs in a reliable fashion for extended periods of time. In recent work we have presented why this is useful [15, 16]. Therefore, it is necessary to develop tools that also allow the continuous control of external parameters of cell culturing, i.e. an incubator is needed.

Many low-cost and open-source incubators can be found in the literature [17, 18]. Some of these incubators focus on achieving lower oxygen concentrations than what is present in ambient air, as such conditions can aid cell growth and viability under certain conditions [19]. These conditions are more physiological and can either be created passively by mixing O_2_ and N_2_ in an open-loop fashion [20], or by actively controlling the corresponding gas concentrations [21]. The latter work has also suggested to flexibly adapt the incubation chamber. However, they have not yet combined this approach with 3D printing technology. Smaller, more portable incubation systems also exist, that can control CO_2_, O_2_, and temperature [22]. Another version is capable of controlling CO_2_ and temperature by repurposing a 3D printer [23]. The CO_2_ concentration was controlled using dry ice. Such approaches, however, still require external power and are not easily adaptable to constraints imposed on the incubation system by experiments. A portable battery-powered incubator exists[24], however, since it is a commercial solution and hence not open-source, it lacks flexibility.

Naturally, incubator solutions are also merged with other lab equipment such as microscopes. Scoggin *et al*. have proposed to 3D print a cell culture dish warming system that can be used during microscopy sessions [25], while Walzik *et al*. have published an open-hardware combination of an incubator and a microscope [26]. They can actively control the CO_2_ concentration and the temperature, as well as passively control the humidity.

Here we present a modular, low-cost, and versatile incubation system, that can be customized to meet a wide range of experimental needs. The majority of the components can be either 3D-printed or easily purchased at electronic shops at a low cost. Our system comprises a central main unit that can be connected to an interchangeable incubation chamber. The main unit has a simple user interface with a display, showing the current temperature, CO_2_ concentration, and humidity present in the incubation chamber. Three push buttons allow to control the heating state (ON/OFF), the mode (Hot/Cold, when used with a Peltier element) and the presence of a stabilized CO_2_ environment (ON/OFF). Two rocker switches allow to turn the power of the system on as well as to connect the battery. A slide switch provides a locking mechanism to avoid accidental actuation of any control buttons.

We introduce two distinct incubation chambers illustrating the versatility of our platform. The combination of the main unit with the “inkugo” incubation chamber yields a compact, portable incubator, allowing for the transportation of living cells for up to two hours without external power. During transport, the incubator is powered by a lithium-ion battery. In addition, the CO_2_ level of the incubator can be held stable at 5 % without an external CO_2_ source by using a 16 g CO_2_ cartridge. The inkugo system was used in previous publications to transport cells from a campus with a cell culture facility to a campus with a scanning electron microscope (SEM) [27].

In addition, we introduce an incubation chamber called “inkubox”, which serves as a system for electrophysiological recordings from neural cultures over extended periods of time. Four recording units (MaxOne, Maxwell Biosystems, Switzerland) for complementary metal–oxide–semiconductor (CMOS) microelectrode arrays (MEA) can be placed within the inkubox chamber for parallel recordings of four neural cultures in a stable environment. Such CMOS MEAs provide an electrode pitch of 17.5 *μ*m and hence enable the recording of extracellular activity in neural cultures with subcellular resolution. We used the here presented technology in previous work for continuous recordings from engineered neural networks on high-density MEAs *in vitro* [15, 16].

The versatility of the presented system is boundless and the incubation chamber can be adapted to meet different experimental needs. For example, employing an H-bridge on the printed circuit board (PCB) allows to drive a Peltier element. This can be used to cool or heat biological samples in a single device.

In summary, the system can be utilized by researchers to introduce an environment compatible with cell culturing to any equipment that previously lacked that capability. The devices shown in this work are cost-efficient incubators that can be used in stationary or mobile use cases.

## 2 Hardware description

The uniqueness of our approach stems from the modularity of our system. By splitting the control unit and the incubation chamber into two separate entities, one control unit can be utilized to control a variety of custom incubation chambers. The incubation chamber can be adjusted to match highly specific experimental needs. Additional incubation chambers tailored to application-specific needs can be designed rapidly, as the electronic equipment required per incubation chamber is kept at a minimum.

The lack of special equipment required during construction as well as the repurposing of readily available and cheap day-to-day equipment makes it an easy-to-reproduce system. Most design components can be printed with conventional 3D printers, while necessary aluminum-based design components can be easily fabricated in any workshop or ordered from manufacturers online. With a cost well below $1000, our system is by an order of magnitude cheaper than commercial equivalents for both the mobile and stationary incubation systems.

A schematic of the system is provided in Fig. 1. A control unit comprises the energy storage and delivery to the incubation chamber, CO_2_ supply, and presents the state of the incubation chamber on a display. Energy to drive a heating element or a Peltier element is provided either via a 13 V DC input or via an integrated lithium-ion battery that can simultaneously be charged when the control unit is plugged in. CO_2_ is provided by a 16 g CO_2_ cartridge, commonly used to inflate bike tires. The cartridges can be inserted into a miniature pressure regulator, commonly used for carbonation of homebrew beer. The manometer present in the pressure regulator can be used to gauge when the cartridge needs to be replaced. The mainboard within the control unit can be connected by an 8-pin Molex cable to a peripheral board, which is placed within the incubation chamber. Moreover, a 6 mm tube connecting the main unit and the incubation chamber allows the flow of CO_2_ from the control unit to the incubation chamber.

**Figure 1.**
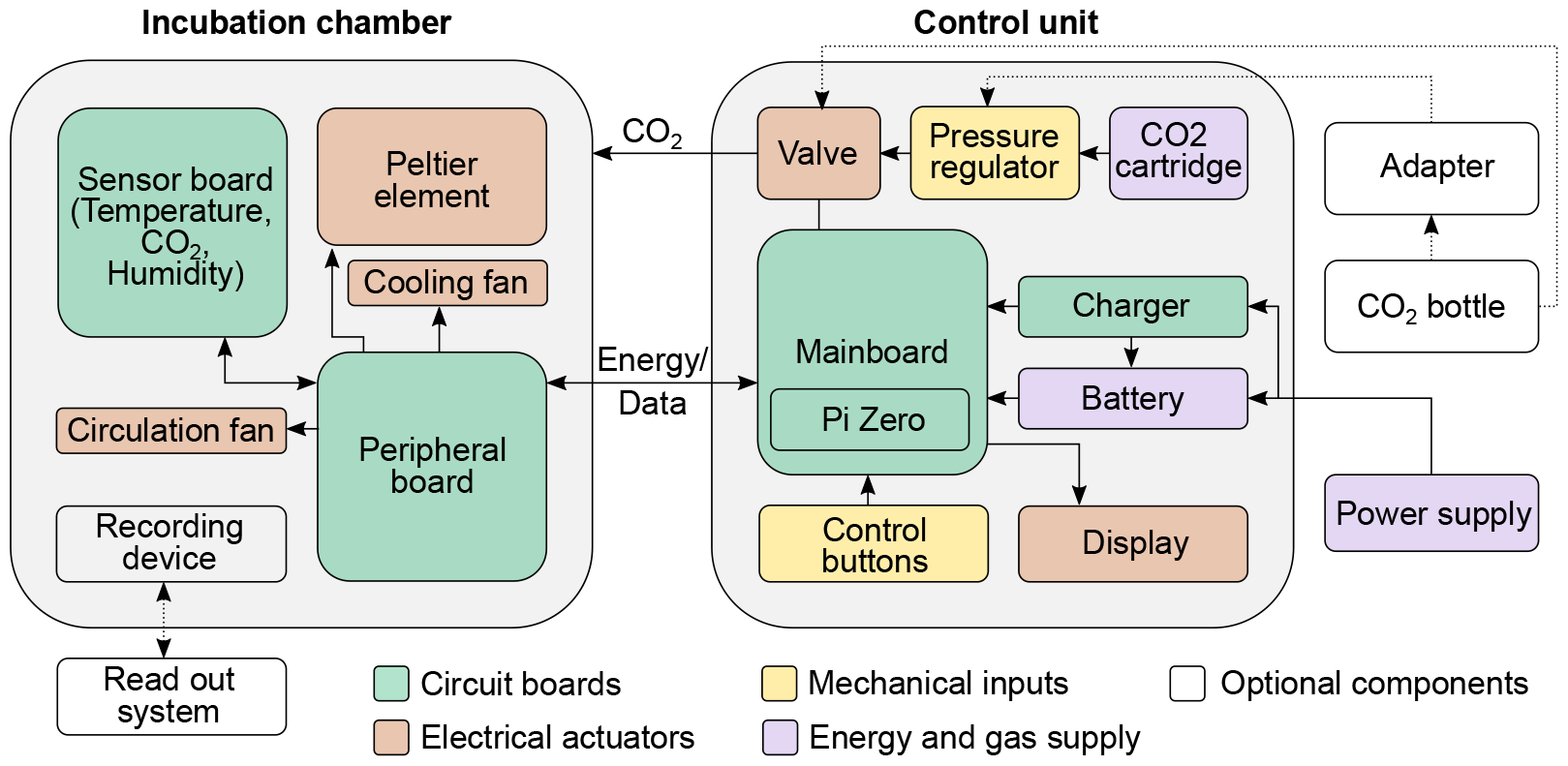
Schematic of the incubator system. The incubator system consists of two separate units: a control unit and an incubation chamber. The two units are connected by a tube for CO_2_ infusion and a cable for heating and data transmission. CO_2_ can be supplied to the control unit by a CO_2_ cartridge or an external CO_2_ bottle. The incubation chamber can be adapted to several experimental needs without requiring a change in the control unit.

An overview of the system with the inkugo configuration is shown in Fig. 2. The incubation chamber can either be placed next to the main unit or the entities can be stacked on top of each other. With a total weight of approximately 3 kg and dimensions of 20 *×* 34 cm, inkugo is suitable for transportation. The incubation chamber measures 12 *×* 12 *×* 9 cm, which provides enough space to transport four stacked Petri dishes (10 cm diameter, 2 cm height). Inkugo can assist researchers and enhance collaboration in decentralized laboratory settings by allowing the exchange and transport of living cell cultures.

**Figure 2.**
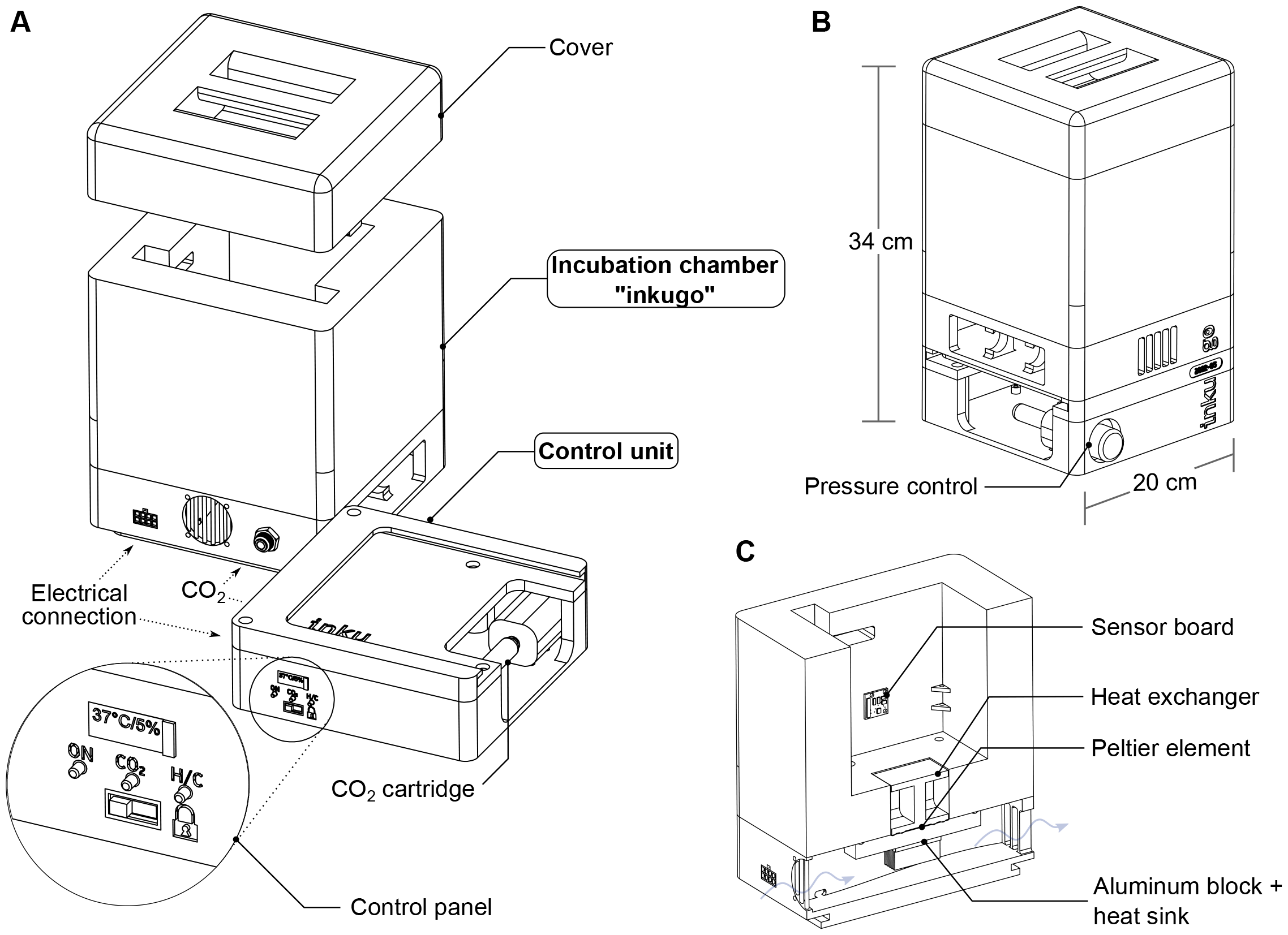
Overview of the inkugo system intended for mobile use. In this use case, the incubator system is comprised of a control unit and a small, lightweight incubation chamber called “inkugo”. **A** The control unit contains a control panel with a display presenting the current state of the culture environment (temperature, CO_2_, and humidity) and buttons to control the device. **B** The inkugo chamber can be stacked on top of the main unit for the transportation of the whole system. **C** Interior of the incubation chamber. A sensor board for temperature, CO_2_, and humidity measurement is placed on the wall of the chamber. A Peltier element heats (or cools) the interior of the chamber. The heat produced by the Peltier element is conducted via an aluminum block and a heat sink.

An overview of the inkubox system configuration is shown in Fig. 3. We have designed the inkubox chamber such that four recording units can be placed inside to enable continuous long-term recording from CMOS MEAs with neural cultures under physiological conditions. The inkubox chamber is split into two compartments measuring 29 cm *×* 18 cm *×* 4.5 cm and 29 cm *×* 10 cm *×* 4.5 cm. A drawer within the inkubox chamber serves as a water bath that can be filled with deionized water to enhance humidity and hence mitigate the evaporation of the cell culture medium. CO_2_ can be supplied by either an external CO_2_ bottle or by a 16 g CO_2_ cartridge integrated into the control unit. Inkubox allows every laboratory with an interest in electrophysiology to continuously record from neural cultures in addition to establishing a cheap incubation system for stationary use. Due to the comparatively large size of inkubox, not every out-of-the-box 3D printer can be used to print it. This work has used an Ender 5 Plus (Shenzhen Creality 3D Technology Co Ltd, China) 3D printer. However, due to the flexibility of the design, a smaller box can also be created.

**Figure 3.**
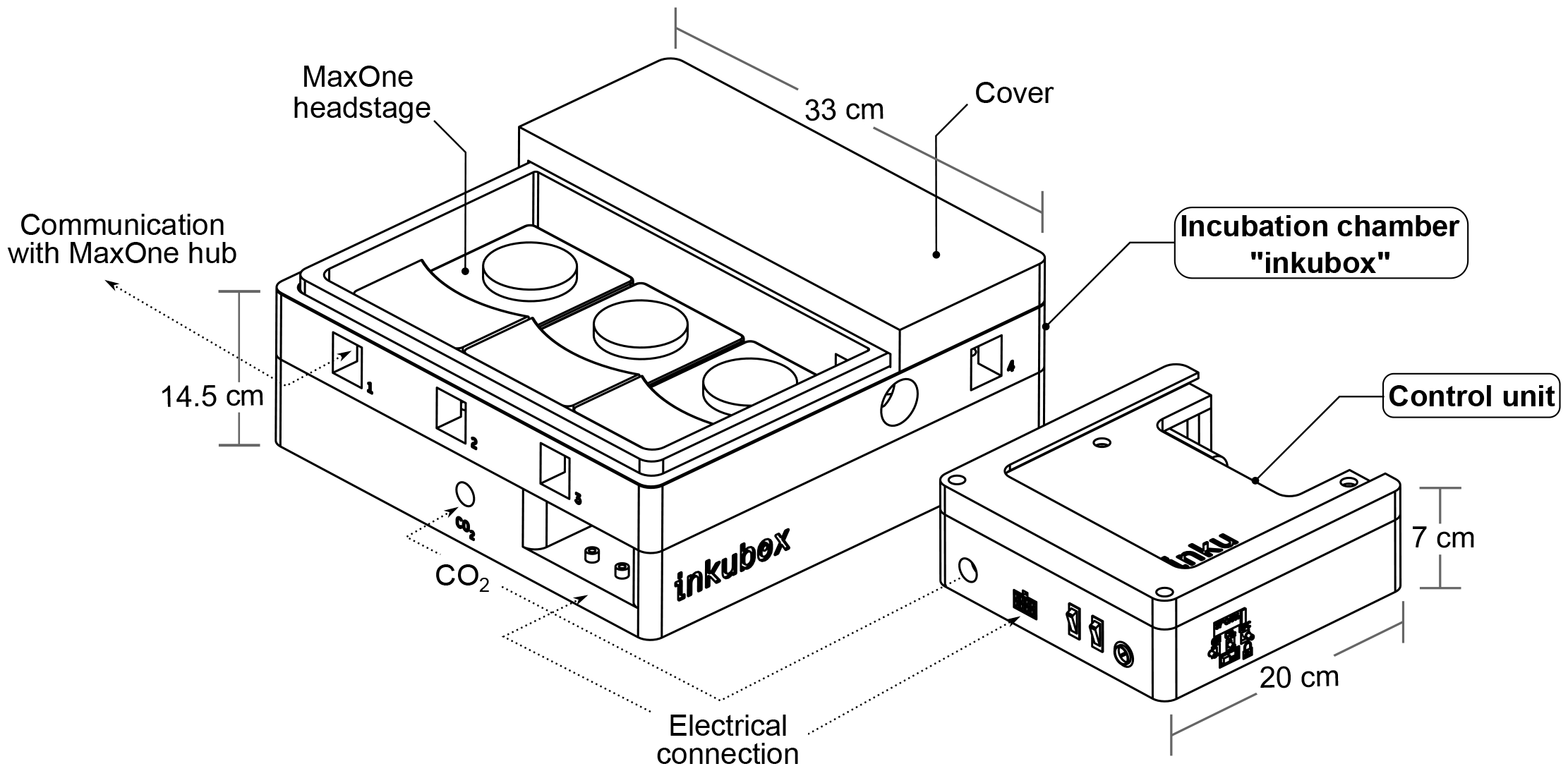
Overview of the inkubox system intended for long-term electrophysiological experiments. Here, the incubator system is comprised of a control unit and an incubation chamber called “inkubox”. The incubation chamber provides space for four MaxOne headstages used to obtain neural recordings from CMOS MEAs. The headstages are connected to MaxOne hubs (recording units) via LAN connectors through the inkubox wall, allowing the hubs to be stationed outside of the controlled incubation environment.

Photos of a fully assembled inkugo and inkubox system are shown in Fig. 4 and Fig. 5, respectively. In summary, our platform provides the following key advantages:

**Figure 4.**
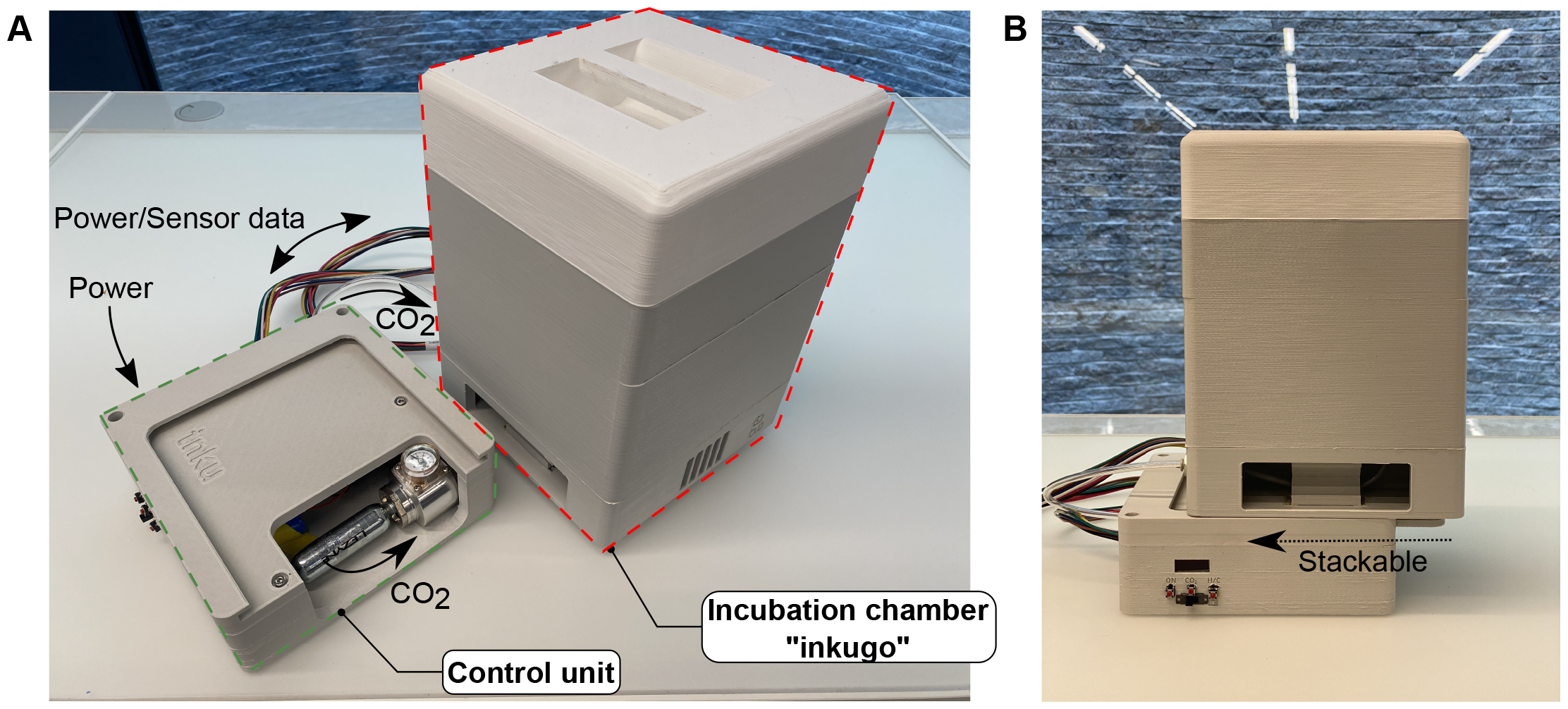
Fully assembled inkugo. **A** Incubation chamber and control unit placed next to each other and connected by a tube and a power/data cable. The CO_2_ source is a threaded 16 g CO_2_ cartridge. **B** Incubation chamber and control unit are stacked on top of each other, simplifying transportation.

**Figure 5.**
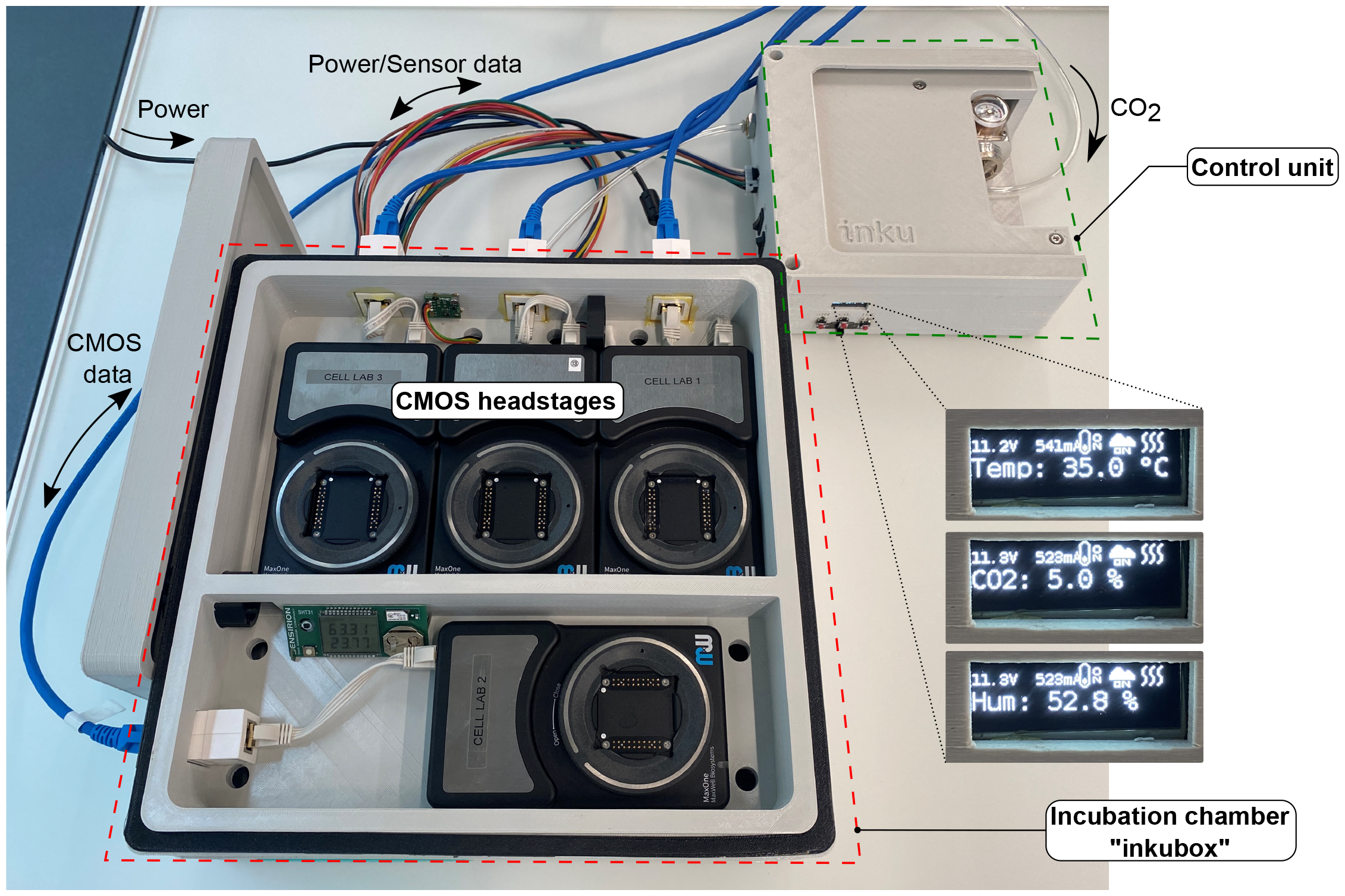
Fully assembled inkubox. The Incubation chamber and control unit are placed next to each other and connected by a tube and a power/data cable. The covers of the incubation chamber were removed to show the recording units placed within. Data from the MEAs are streamed to a recording PC via LAN cables. When used in the here presented stationary mode, the valve (within the control unit) can by connected directly to an external CO_2_ bottle.

- an easy-to-manufacture system using 3D printing and basic metal working
- a simple user interface and easy operation with just a few buttons
- a unitized control software written in Python that can be adapted and extended easily
- a highly modular system allowing to be adapted to a wide range of experimental needs and cell types at variable target temperatures and CO_2_ concentrations.
- a fully-designed incubation chamber (inkugo) to transport cells for more than 2 hours, encouraging collaboration in decentralized lab-settings
- a fully-designed incubation chamber (inkubox) to be used in stationary mode for long-term recordings from neural cultures

## 3 Design files summary

All files are published under a CC BY-SA 4.0 open source license and can be found online. The design files were given unique designators (1A - 3H) which are mentioned both in the build instructions and the corresponding figures (Fig. 6, 7, 8, and 9). The control unit design files are comprised of a cover (1A) and a bottom enclosure (1B) that the mainboard, battery, valve, and pressure controller can be attached to. Both files (1A, 1B) need to be 3D-printed using PLA.

**Figure 6.**
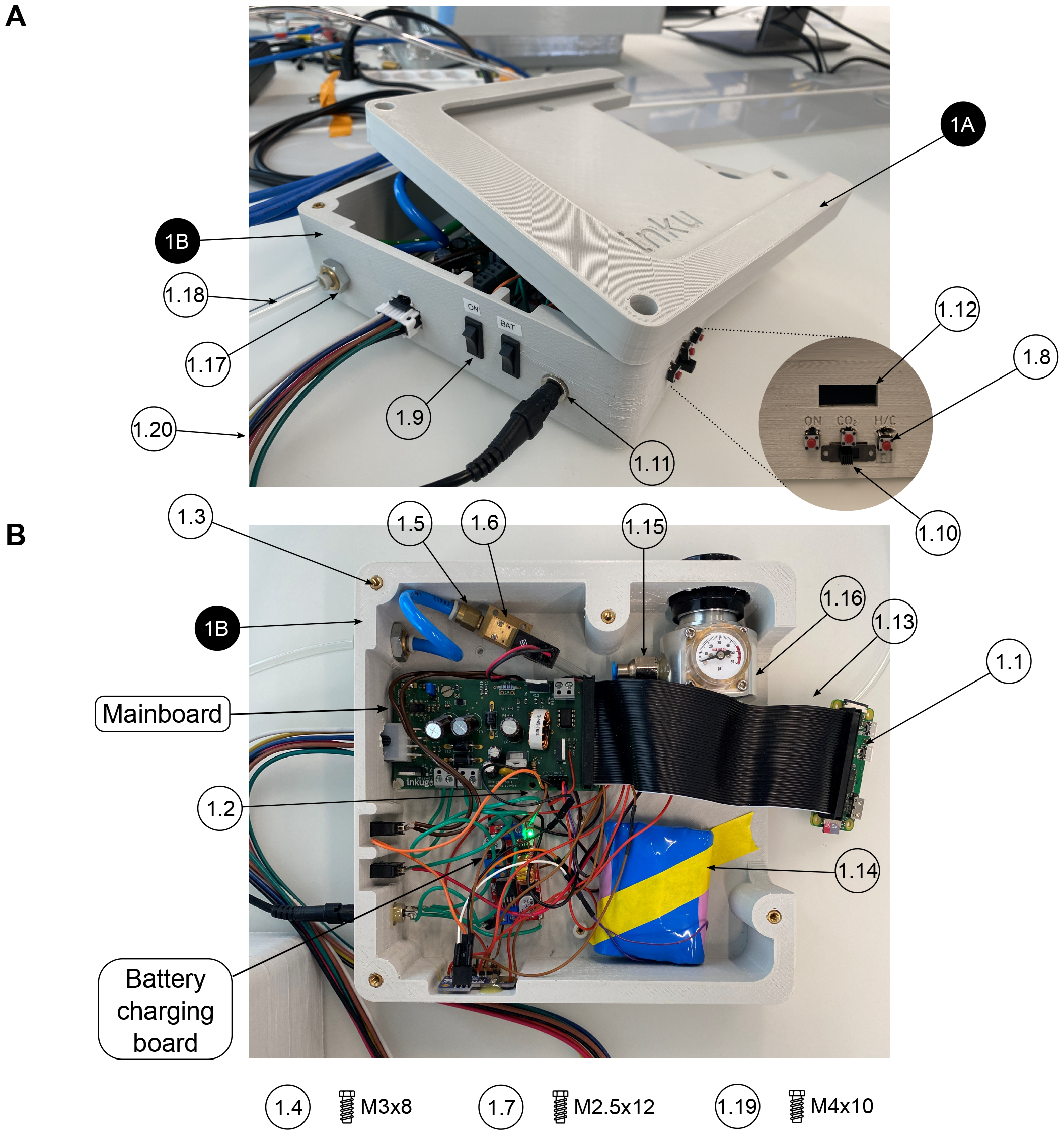
Assembly of the control unit. Design files are indicated with a black pin, other components are shown with a white pin. All components are listed in the bill of materials in the supplementary information. **A** Exterior view of the control unit with the cover partially removed. **B** Top view of the interior of the control unit with removed cover.

**Figure 7.**
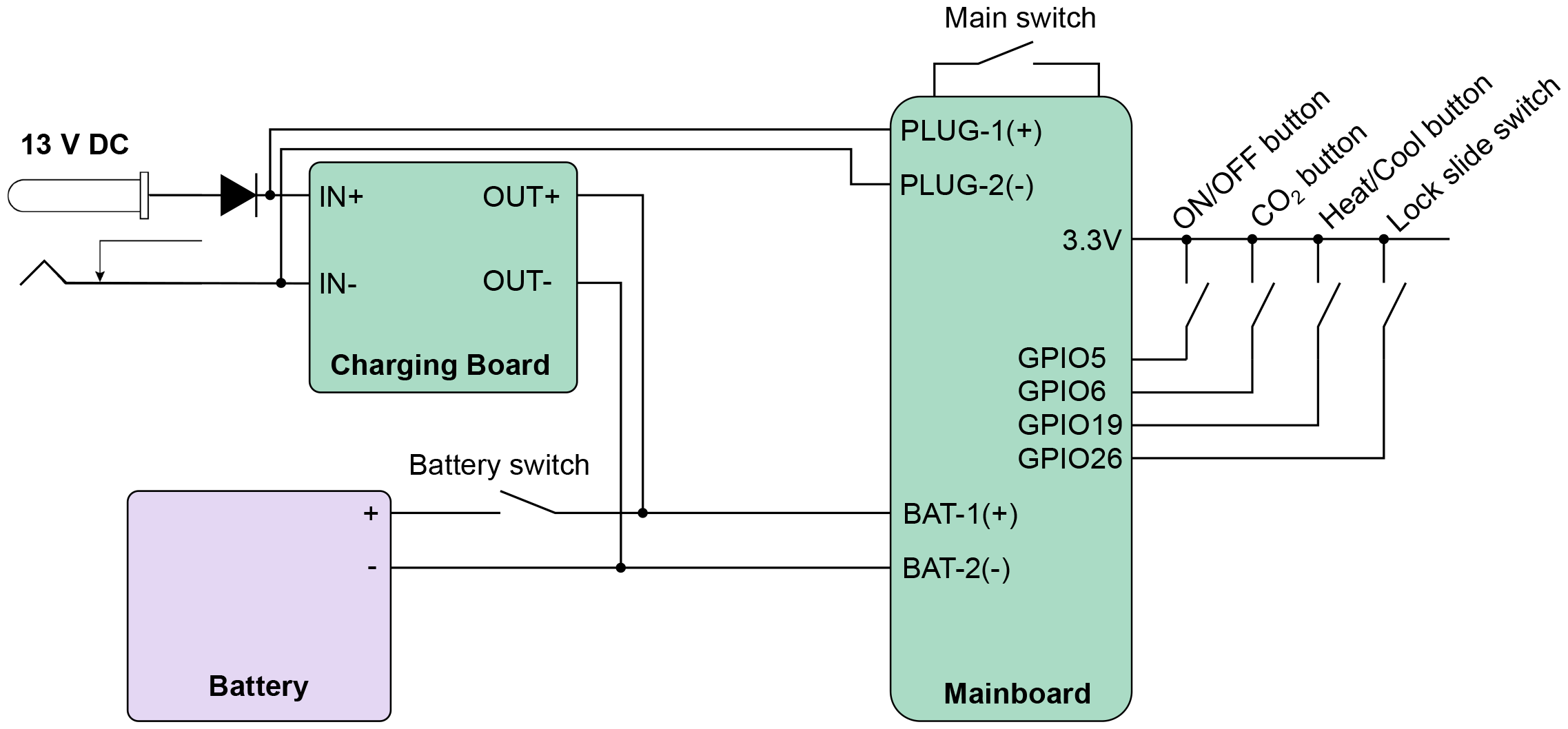
Wiring diagram of the power and control circuit of the control unit. The incubator system is powered with a 13 V DC power supply or a lithium-ion battery, that is charged via a charging board. The battery allows the incubation system to be disconnected from an external power source making this incubation solution portable. The battery can be connected to the charging board via a switch (“Battery switch”). The control unit can be turned on with another switch (“Main switch”). Three push buttons control the power state, whether CO_2_ should be supplied to the incubation chamber, and the polarity of the applied voltage to the Peltier element to switch from heating to cooling. A slide switch provides a locking mechanism to prevent accidental operation of the device.

**Figure 8.**
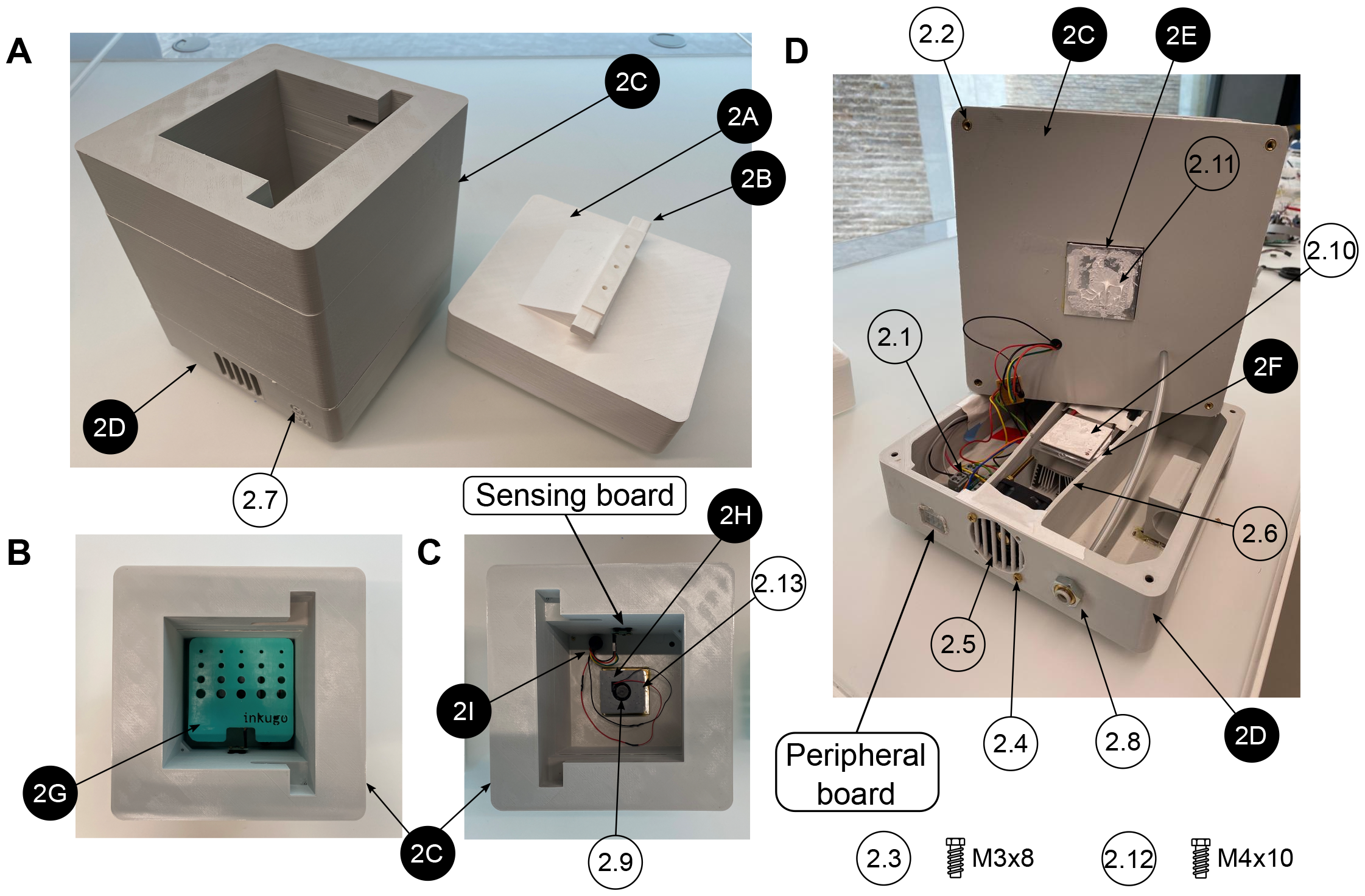
Assembly of the inkugo incubation chamber. Design files are indicated with a black pin, other components are shown with a white pin. All components are listed in the bill of materials in the supplementary information. **A** Side view of the assembled inkugo chamber with removed cover. **B** Top view of the interior of the incubation chamber. **C** Top view of the interior of the incubation chamber without the sample holder. **D** Side view of the disassembled inkugo chamber.

**Figure 9.**
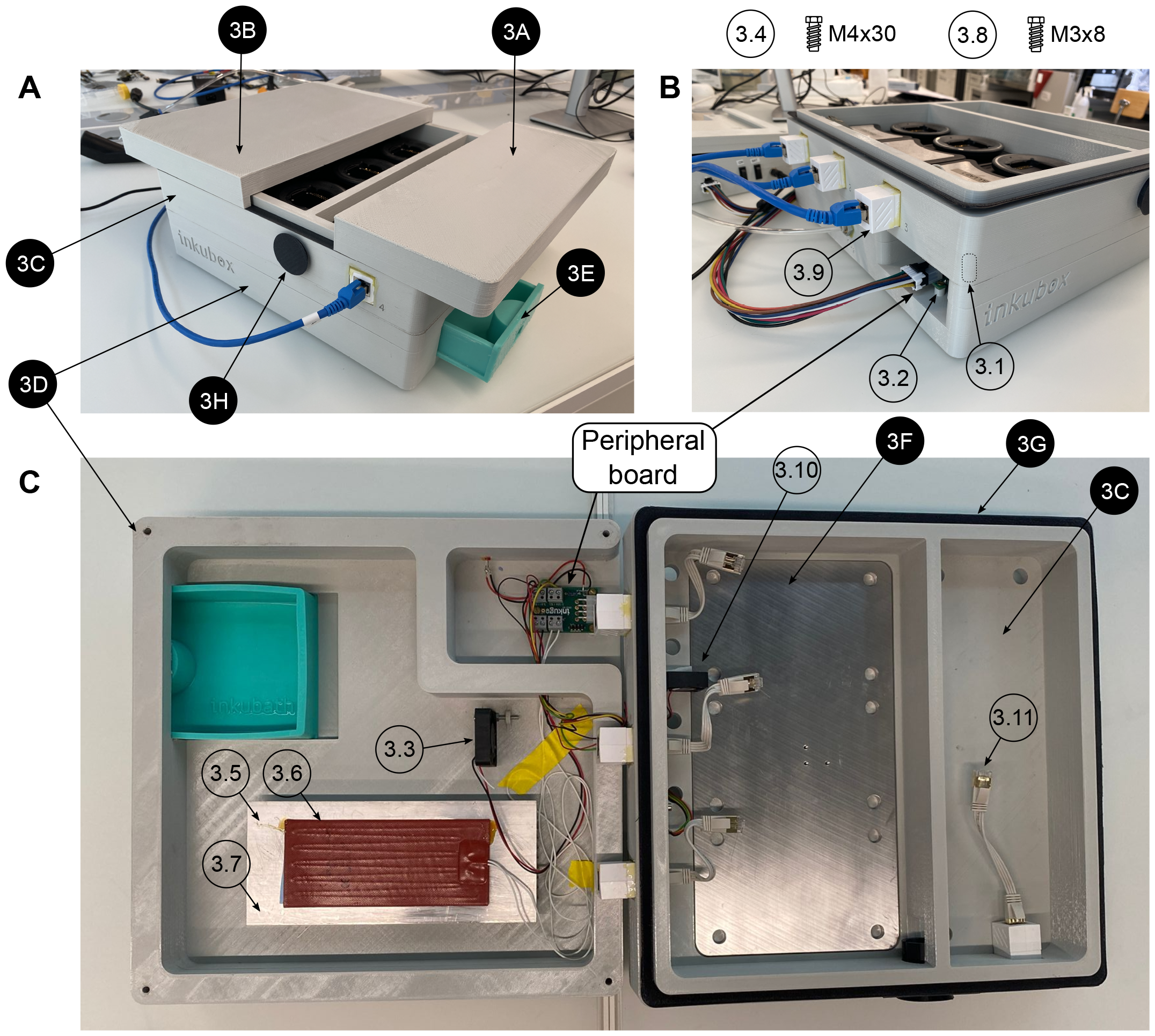
Assembly of the inkubox incubation chamber. All components are listed in the bill of materials in the supplementary information. Design files are indicated with a black pin, other components are shown with a white pin. **A** Side view with covers (3A and 3B) slightly shifted. **B** Side view illustrating the gas, electrical, and data connections to and from the inkubox. **C** Top view of the disassembled inkubox chamber.

The inkugo (Fig. 4 and 8) incubation chamber design files consist of a cover (2A, PLA), a connection bar (2B, PLA) to be screwed on the cover, the incubation chamber center part (2C, PLA) for cell storage and housing the sensor board, and the bottom part of the incubation chamber (2D, PLA) that holds the peripheral board and ventilation components. In the fully printed design, the cover (2A) can be interlocked with the center part (2C) via the connection bar (2B). The connection bar (2B) can be adjusted to control the tightness of the fit. Moreover, an aluminum block (2E) is needed to conduct heat from the Peltier element into or out of the chamber. The Peltier element is placed on top of an aluminum plate (2F), on which three heat sinks can be attached. Biological samples can be placed onto the sample holder (2G, resin). Air circulation inside the chamber is achieved through a small fan that is placed inside the fan holder (2H, PLA).

The inkubox (Fig. 5 and 9) design files consist of a small incubation chamber cover (3A, PLA) and a larger chamber cover (3B, PLA), which can be placed separately on top of the main chamber (3C, PLA). Having two separate chambers (a small one with one recording unit and a large one with three recording units) allows to access the recording units independently. This can for example protect one MEA from light exposure when operating the other three headstages. The main chamber (3C) is screwed onto a bottom chamber (3D), which is housing a heater, a circulation fan, the peripheral board, and a water bath (3E, resin). An aluminum plate (3F) serves as a heatsink for three headstages to conduct excess heat that can arise from any electrical equipment introduced to the incubator. The insulation ring (3G, TPC Flex) can be placed in between the main chamber (3C) and the covers (3A, 3B). Moreover, the main chamber (3C) has an opening on the side that makes it possible to insert cables into the main chamber. This could for example serve as an entry site for a LEMO-BNC cable to connect a headstage to an external device or for different stimulation modalities such as light. The flexible plug (3H) allows for such cable insertions while isolating the main chamber well enough to hold the temperature and CO_2_ concentration stable.

|  | Design filename | File type | Location of the file |
| --- | --- | --- | --- |
| 1A | 1A_control_unit_top | 3D File | ./01_Mechanical/Control_unit |
| 1B | 1B_control_unit_bottom | 3D File | ./01_Mechanical/Control_unit |
| 2A | 2A_incubation_chamber_cover | 3D File | ./01_Mechanical/inkugo |
| 2B | 2B_incubation_chamber_connector | 3D File | ./01_Mechanical/inkugo |
| 2C | 2C_incubation_chamber_center | 3D File | ./01_Mechanical/inkugo |
| 2D | 2D_incubation_chamber_bottom | 3D File | ./01_Mechanical/inkugo |
| 2E | 2E_incubation_chamber_aluminum_block | 3D File +<br>CAD | ./01_Mechanical/inkugo |
| 2F | 2F_incubation_chamber_heat_sink_plate | 3D File +<br>CAD | ./01_Mechanical/inkugo |
| 2G | 2G_inubation_chamber_sample_holder | 3D File | ./01_Mechanical/inkugo |
| 2H | 2H_incubation_chamber_small_fan_holder | 3D File | ./01_Mechanical/inkugo |
| 2I | 2I_incubation_chamber_cable_plug | 3D File | ./01_Mechanical/inkugo |
| 3A | 3A_small_cover | 3D File | ./01_Mechanical/inkubox |
| 3B | 3B_large_cover | 3D File | ./01_Mechanical/inkubox |
| 3C | 3C_main_chamber | 3D File | ./01_Mechanical/inkubox |
| 3D | 3D_bottom_chamber | 3D File | ./01_Mechanical/inkubox |
| 3E | 3E_water_bath | 3D File | ./01_Mechanical/inkubox |
| 3F | 3F_metal_plate | 3D File +<br>CAD | ./01_Mechanical/inkubox |
| 3G | 3G_insulation_ring | 3D File | ./01_Mechanical/inkubox |
| 3H | 3H_flex_plug | 3D File | ./01_Mechanical/inkubox |
|  | Connection_Board | PCB Schemat-<br>ics | ./02_Electrical/Connection_Board |
|  | Mainboard | PCB Schemat-<br>ics | ./02_Electrical/Mainboard |
|  | Sensor_Board | PCB Schemat-<br>ics | ./02_Electrical/Sensor_Board |
|  | Code | Python code +<br>Operating sys-<br>tem | ./03_Code |
|  | Printing Profiles | Configuration<br>Files | ./04_Printing_Profiles |

### Bill of materials

The complete bill of materials can be found in the supplementary information. The electrical components for all PCBs are listed in a separate spreadsheet that can also be found in the supplementary information. We have used the following equipment to manufacture the devices:

- soldering iron and reflow oven
- LCD resin printer (Sonic Mini 4K, Phrozen, Taiwan)
- extrusion printer (Ender 5 Plus, Shenzhen Creality 3D Technology Co Ltd, China)
- epoxy glue (used here: Araldite Rapid, Huntsman Corporation, USA)
- screwdrivers

CO_2_ cartridges are listed separately since they serve as single-use consumables. Cartridges of 16 g or 25 g CO_2_ can be purchased for less than $1/cartridge from many bike shops.

## 4 Build instructions

1. 3D-print all parts for the control unit and the desired incubation chamber (inkugo or inkubox).
2. Solder all components onto the three PCBs (mainboard, connection board, and sensor board). For soldering SMD components, a reflow oven is recommended.
3. Set up the Raspberry Pi Zero (#1.1) using the SD card image, which is provided in the design files repository (Folder “03 Code”). Alternatively, set up the Raspberry Pi with another operating system and run on it the code provided on github^1^.
4. Using a laboratory power supply, set the constant current (CC) and constant voltage (CV) limits of the battery charging board (using the battery listed in the bill of materials, the settings used were V_*CV*_ = 12.6 V and I_*CC*_ = 1 A).
5. Assembly of the control unit (see Fig. 6 and 7):
  a. Using a soldering iron, place the M3 (#1.2) and M4 (#1.3) thread inserts into the 3D-printed openings.
  b. Fix the mainboard to the main unit bottom piece (1B) using three M3x8 screws (#1.4).
  c. Connect two M5-6.0 mm fittings (#1.5) to the valve (#1.6).
  d. Fix the valve to piece 1B using two M2.5x12 screws (#1.7).
  e. Using epoxy glue, glue the three push buttons (#1.8), two rocker switches (#1.9), one slide switch (#1.10), the DC power jack (#1.11), and the display (#1.12) to the exterior of piece 1B.
  f. Connect the Raspberry Pi (#1.1) to the mainboard using the GPIO ribbon cable (#1.13).
  g. Wire the switches and buttons to the mainboard and battery according to the schematics shown in Fig. 7.
  h. Connect the battery (#1.14) to the mainboard and the battery charging board according to Fig. 7.
  i. Connect the M8x0.75-6 fitting (#1.15) to the output of the CO_2_ mini pressure regulator (#1.16).
  j. Mount the CO_2_ mini pressure regulator to piece 1B using four M3x8 screws (#1.4).
  k. Using two nuts, fix the wall mount (#1.17) for the CO_2_ tube to piece 1B.
  l. Connect the pressure regulator to the valve and the valve to the wall mount using a 6 mm tube (#1.18) of arbitrary length.
  m. Solder the push buttons (#1.8), rocker switches (#1.9), and the DC power jack (#1.11) to the main and battery charging board according to the schematics shown in Fig. 7.
  n. Connect the display (#1.12) to the mainboard using the display connect pins (SDA/SCL/3.3V/GND) labeled “DP CONNECT”.
  o. Screw the main unit cover (piece 1A) to piece 1B using four M4x8 screws (#1.19) to finish the assembly of the main unit.
6. Assembly of the incubation chamber:
  a. inkugo chamber (see Fig. 8):
    1. Using a soldering iron, place all M3 and M4 thread inserts (#2.1 and #2.2) into the inkugo chamber pieces (2A-2D).
    2. Fix the peripheral board to the bottom piece of the incubation chamber (piece 2D) using three M3x8 screws (#2.3).
    3. Using M4x30 rods (#2.4), fix the Peltier element cooling fan (2.5) to piece 2D in the central channel.
    4. Mount the three heatsinks (#2.6) to the heatsink plate (piece 2F).
    5. Place the fully assembled heatsink plate inside of the central channel in piece (2D) next to the cooling fan (2.5) so that the fan provides airflow through the heatsinks.
    6. Connect the Peltier element cooling fan (2.5) to the appropriate pins on the peripheral board (12V/F GND).
    7. Glue a 3 mm LED (#2.7) to the opening in the inkugo logo on the side of piece 2D as a “POWER OK” indicator and connect the LED to the peripheral board (pads labeled “LED”).
    8. Fix the wall mount (#2.8) for the CO_2_ tube to piece 2D using two nuts.
    9. Screw the sensing board to piece 2C using M3x8 screws (#2.3).
    10. Connect the sensing board (SCL/SDA/3.3V/GND) to the peripheral board (3.3V/GND).
    11. Place the chamber fan (#2.9) inside the fan holder (piece 2H) and connect it to the peripheral board (3.3V/GND).
    12. Using epoxy glue, glue the heat exchanger (piece 2E) to the bottom square opening of the centerpiece of the incubation chamber (piece 2C).
    13. Place the Peltier element (#2.10) between the heat exchanger (piece 2E) and the heat sink plate (piece 2F) using thermal paste (#2.11) to enhance thermal conductivity.
    14. Connect the Peltier element (#2.10) to the peripheral board.
    15. Screw pieces 2C and 2D together using M4x10 screws (#2.12).
    16. Using thermal paste (#2.11), add the planar heatsink (#2.13) onto piece 2E inside the incubation chamber.
    17. Place the chamber fan (#2.9) and fan holder (piece 2H) on top of the heatsink (#2.13).
    18. Connect the incubation chamber cover (piece 2A) to the chamber connector (piece 2B) using three M3x8 screws.
    19. Close the wire hole inside the incubation chamber using the cable plug (piece 2I)
    20. Place the sample holder (piece 2G) inside the incubation chamber.
  b. inkubox chamber (see Fig. 9):
    1. Using a soldering iron, place four M4 thread inserts (#3.1) into the main chamber piece (3C) and three M3 thread inserts (#3.2) into the bottom chamber (3D).
    2. Screw the peripheral board onto the bottom chamber (piece 3D).
    3. Fix the circulation fan (#3.3) to piece 3D using an M4x30 screw (#3.4).
    4. Using thermal paste (#3.5), place the silicon heat foil (#3.6) onto the large heatsink (#3.7).
    5. Fix the heatsink (#3.7) to piece 3D using epoxy glue.
    6. Connect the heat foil (#3.6) to the peripheral board.
    7. Connect the sensing board to the centerpiece of the incubation chamber (piece 3C) using three M3x8 screws (#3.8) and connect the sensing board to the peripheral board (SDA/SCL/3.3V/GND).
    8. Using epoxy, glue the four LAN connectors (#3.9) in the four rectangular openings of piece 3C.
    9. Glue the chamber fan (#3.10) to piece 3C outside the opening of the metal plate perpendicular to the wall of piece 3C.
    10. Screw pieces 3C and 3D together using four M4x30 screws (#3.4).
    11. Place the metal plate (piece 3F) in the opening of piece 3C.
    12. Glue the insulation ring (piece 3G) on top of piece 3C using epoxy.
    13. Close the incubation chamber with covers 3A and 3B.
    14. Insert the 3D-printed water bath (piece 3E) into the designated opening in piece 3D.
    15. Close the opening on the side of piece 3C with the flexible plug (piece 3H).
    16. Insert the short LAN cables (#3.11) into the inside-facing opening of the LAN wall connectors (#3.9).
7. Connect the main unit to the incubation chamber using the 8-pin Molex cable (#1.20).
8. Connect the main unit and the incubation chamber using a 6 mm tube (#1.18) and connect the main unit to a CO_2_ source (gas cartridge or gas bottle).

## 5 Operation instructions

1. Connect the incubation chamber to the control unit (8-pin cable and gas tube).
2. Connect the control unit to a power outlet using the DC charger.
3. Connect the control unit to a CO_2_ source. When using an external CO_2_ bottle, connect the tube directly to the valve (the incoming pressure should not exceed 2 bar). When using a 16 g CO_2_ cartridge, screw it into the opening of the pressure controller, while the controller is set to “OFF”.
4. Open the controller by rotating the knob to “LOW”.
5. Switch on the control unit with the “ON” rocker switch.
6. Wait until the device boots up.
7. Click the “ON” push button below the display and the heating begins. For cooling, the system has to be set to “Cool”. For cooling the display will show a small snowflake icon. When the system is in heating mode, it will show a little flame instead.
8. When the system reaches its target temperature (default is 37 °C), calibrate the CO_2_ sensor. For this, press the “CO_2_” button for at least 5 s and release. A message will be shown on the display confirming that the calibration was successful. The CO_2_ concentration inside of the chamber needs to be at 0 % during the calibration.
9. Press the “CO_2_” button once to start the inflow of “CO_2_” into the incubation chamber.
10. When the “CO_2_” level reached its target concentration (default is 5 %), place the cells inside the incubation chamber.

## 6 Validation and characterization

The incubation system was characterized by continuously writing the temperature and CO_2_ values into a log file once the system was started. Fig. 10 shows the temperature and CO_2_ level within the inkugo incubation chamber over time with and without active control. After the heating was initiated, the temperature increased to the target temperature of 37 °C within a timeframe of approx. 15 min. After 50 min, the heating process was stopped and the temperature decayed in an exponential manner, reaching the ambient temperature after approximately 90 min. The target CO_2_ concentration was reached in a manner of a few minutes. A single 16 g CO_2_ cartridge was sufficient to operate the inkugo system for more than 24 h, while the battery lasted for approx. 2 h, when the system was already heated up to the target temperature. The discharge curve of the battery is shown in the supplementary information in Fig. S1. Combining the main unit with a highly insulating incubation chamber made out of polystyrene allows the establishment of a system capable of cooling, when set to the cooling mode (Fig. S2). The assembly instructions for this version of inkugo are given in the supplementary information. The temperature curve in this mode is shown in the supplementary information in Fig. S3.

**Figure 10.**
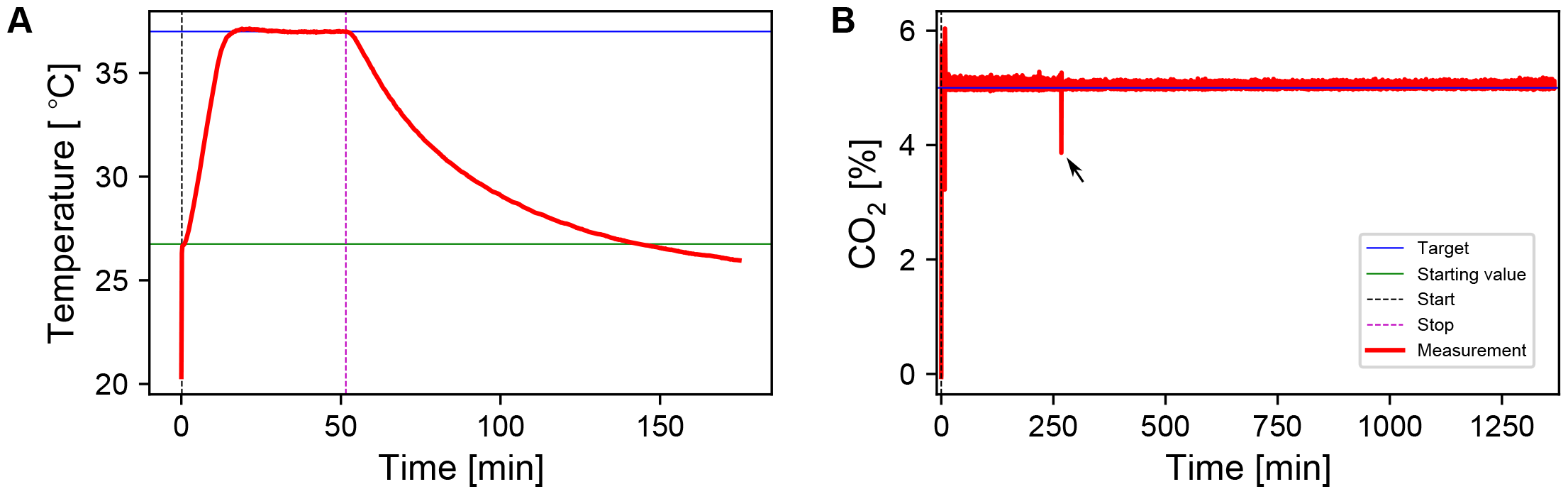
Characterization of the incubator system using the inkugo chamber. **A** Temperature behavior and stability over time. The actual measured values are plotted in red, while the black and purple dashed lines indicate the time when the heat and CO_2_ inflow process was started and stopped. The blue and green lines indicate the target and starting values. **B** Long-term stability of the CO_2_ level within the incubation chamber. The target concentration was reached within 2 min and maintained for over 21 h. The black arrow indicates a transient drop in CO_2_ when the chamber was briefly opened to remove a sample.

The inkubox behavior is illustrated in Fig. 11. After a transient phase of approximately 2.5h, the system reaches its target temperature of 37°C. The temperature decay that occurs after stopping the heating process is much slower in contrast to the inkugo system, likely due to the higher heat capacity to heat conductance ratio of inkubox with respect to inkugo. The CO_2_ target of 5 % is reached within a 2 min window. An exponential decay is observable when the CO_2_ injection is stopped. The exponential decay constant of the CO_2_ decay is approximately 0.1 min^−1^.

**Figure 11.**
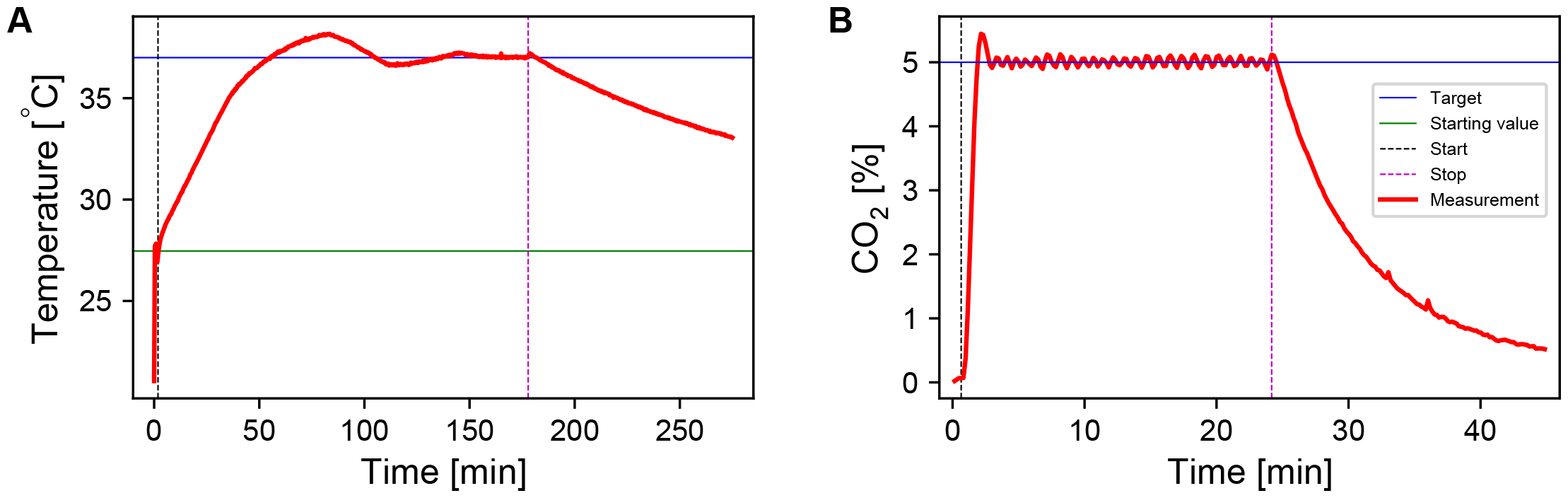
Characterization of the incubation system using the inkubox chamber. **A** Temperature behavior and stability over time. **B** Stability of the CO_2_ level within the incubation chamber over time. The actual measured values are plotted in red, while the black and purple dashed lines indicate the time when the heat/CO_2_ process was started and stopped. The blue and green lines indicate the target and starting values.

To validate inkugo, neural recordings were conducted on an engineered neural network at 84 days *in vitro* (DIV), which was growing on a CMOS MEA inside a PDMS microstructure. The details on cell culturing can be found in the supplementary information (Section: “Cell culturing”).

The MEA was removed from a commercial stationary incubator and placed inside the inkugo chamber, which was preheated to 37°C and reached a CO_2_ concentration of 5 %. After 4.5 h, the MEA was removed from the inkugo chamber and a 1 min electrical recording was performed. As shown in Fig. 12, the neural culture remained spontaneously active, proving that inkugo was able to sustain a physiological environment and keep the culture alive.

**Figure 12.**
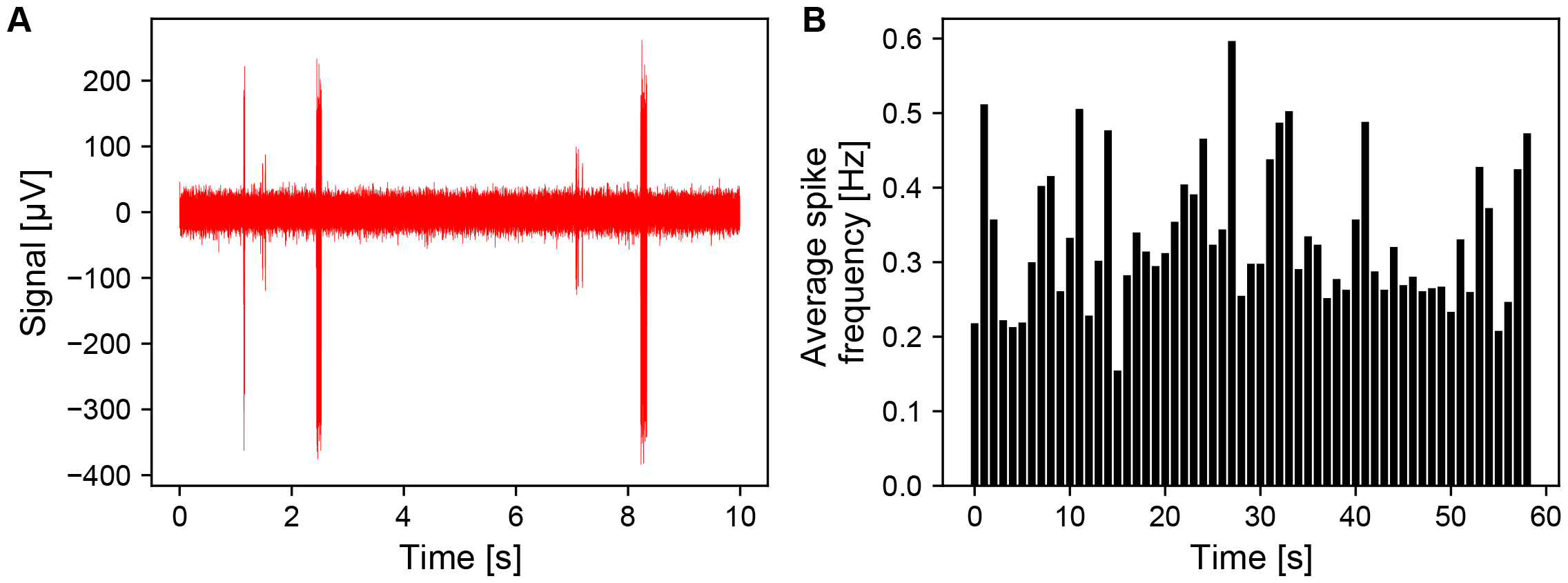
Validation of the inkugo system. Neural recordings were performed on a culture growing on a CMOS MEA at DIV 84. The culture was taken from a commercial stationary cell incubator and placed in inkugo for 4.5 h. Afterwards, spontaneous electrical activity was recorded for 1 min. **A** Spike traces on the most active electrode and **B** average spike frequency per electrode.

Inkubox was validated by performing a spontaneous electrophysiology recording from a neural culture on a CMOS MEA at 80 DIV. The MEA was placed inside a headstage within inkubox. The data of a 6 h recording is shown in Fig. 13. It is evident that the culture remained active throughout the whole experiment. The inkubox system introduces a physiological environment that makes it feasible to record neural culture activity for multiple hours as opposed to only a few minutes without it. The system was used to generate data for previous publications [15, 27]. In these works, neural cultures were maintained in commercial incubators and placed in inkubox several times per week for multiple hours for recording purposes. Neural cultures remained viable and active for months. The lack of active humidity control and automated medium exchange limits the time for continuous recordings. While a detailed analysis would be required, we recommend that cultures may not be placed in inkubox for longer than one day. The system shown in this work provides a cheap and flexible solution to introduce a physiological environment to various experimental settings. However, the system can be further improved to simplify the manufacturing process. For example, the wiring from the buttons/display to the mainboard within the control unit can be simplified: components could be consolidated on another PCB attached to the side of the control unit or fully integrated into the main board. Moreover, the device currently requires the usage of multiple screw types. By reducing the amount of different screws needed, reproducibility is further enhanced. Finally, PLA is both biodegradable and biocompatible [28]. Depending on the use case, one could consider utilizing similarly performing plastics such as polyethylene terephthalate glycol (PETG).

**Figure 13.**
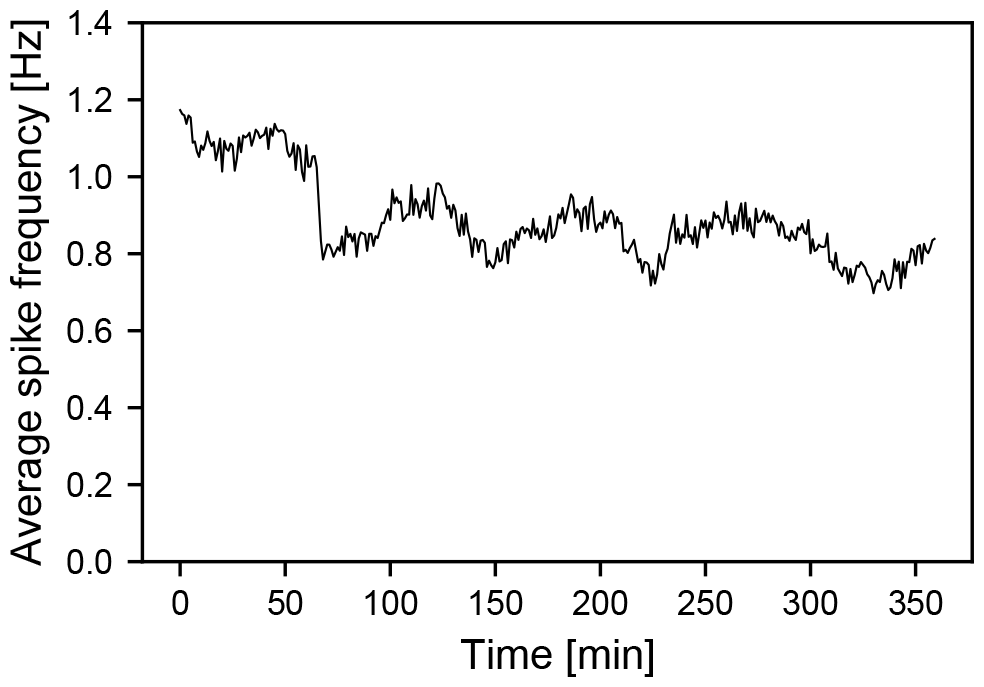
Validation of the inkubox system. A neural culture growing on a CMOS MEA at DIV 80 was placed in inkubox and spontaneous activity was recorded continuously for 6 h.

## 7 Ethics statements

The usage of animal cells was permitted by the veterinary office of the canton Zürich.

## 8 CRediT author statement

**Jens Duru**: Conceptualization, Methodology, Validation, Investigation, Data Curation, Writing - Original Draft. **Benedikt Maurer**: Conceptualization, Methodology, Investigation. **Tobias Ruff** : Conceptualization, Validation, Investigation. **Julian Hengsteler**: Conceptualization, Methodology, Investigation. **Sophie Girardin**: Investigation. **János Vörös**: Conceptualization, Funding acquisition, Project administration. **Stephan J. Ihle**: Conceptualization, Methodology, Validation, Software, Investigation, Data Curation, Writing - Original Draft, Project administration.

## 9 Acknowledgements

Funding: This work was supported by ETH Zürich, Switzerland; the Swiss National Science Foundation [project number 165651]; the Swiss Data Science Center, a FreeNovation grant; and the Human Frontiers Science Program Organization.

## Footnotes

1 https://github.com/lbb-neuron/Inkubox

