## Supplementary Information for "A modular and flexible open source cell incubator system for mobile and stationary use"

Jens Duru, Benedikt Maurer, Tobias Ruff, Julian Hengsteler, Sophie Girardin, János Vörös and Stephan J. Ihle

Laboratory of Biosensors and Bioelectronics, Institute for Biomedical Engineering, Eidgenössische Technische Hochschule (ETH) Zürich, Switzerland

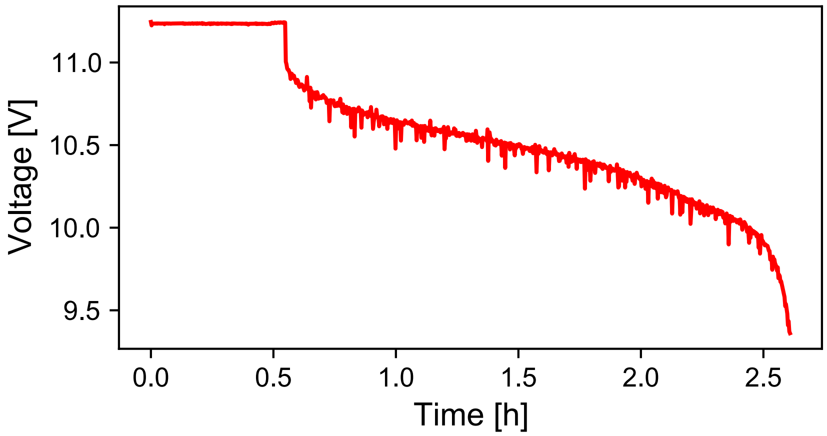


Figure S1: **Battery discharge curve.** The 13 V DC plug was removed from the main unit at approximately 30 min after the system was started. The system was able to run for approximately 2 h powered with the battery.

### 1 Cell culturing

The cells used in this work were primary retinal neurons from E18 rat retinas. The dissociation method was modified from previously established protocols [1]. Cells were seeded into Aggrewell400 plates (Stemcell Technologies, 34415) to form spheroids containing 800 neurons. Spheroids were manually seeded into the seeding wells of PDMS microstructures mounted onto the CMOS array [2]. Spheroids were cultured in RGC medium. 500ml of RGC medium contain 229ml Neurobasal Plus (Gibco, A3582901), 229ml DMEM (Gibco, 11960044), 5ml Glutamax (Gibco, 35050061), 5ml Sodium Pyruvate (100mM, Gibco, 11360070), 5ml Antibiotic-Antimycotic (100X, Gibco, 15240096), 5ml N2 Supplement (100X, Gibco, 17502048), 10ml B27 (50X, Gibco, 17504044), 10ml N21 Supplement (50X, R&D Systems, AR008), 0.5ml NAC Stock (5 mg/mL, Sigma-Aldrich, A8199), 0.5ml Forskolin Stock (4.2 mg/mL, Sigma-Aldrich, F6886), 0.5ml BDNF Stock (50 $\mu$g/mL, Preprotech, 450-02), 0.5ml CNTF Stock (10 $\mu$g/mL, Preprotech, 450-13), 0.5ml NGF 7S Stock (10 $\mu$g/mL, final 10 ng/mL, Preprotech, 450-34), 0.1ml GNDF (10 ng/mL, Preprotech, 450-51). The medium was changed twice per week.

### 2 Polystyrene inkugo

The polystyrene inkugo (p-inkugo) chamber is an adaptation of the main inkugo chamber described in the main text of the manuscript and has the same features. In addition, it can be used to cool down samples by more than 15 K. For this, the current applied to the Peltier element is reversed, allowing the chamber to be cooled down. This feature can also be used when powering p-inkugo with the battery. However, it is not advised, as cooling with Peltier elements is much less energy efficient than heating. Therefore, the discharge of the battery will be considerably faster in comparison to the heating mode.

#### 2.1 Build instructions

The lower part of p-inkugo containing the electronics can be taken over from the main inkugo version or it can be built as described in the “Build instructions” chapter of the manuscript. The following design files are required:

| Designator | Design filename | File type | Location of the file | Comment |
|---|---|---|---|---|
| 4A | 4A_polystyrene_box | N/A | N/A | Polystyrene box of size approx: 240mm x 240mm x 250mm |
| 4B | 4B_sensing_board_holder | CAD | ./01_Mechanical/P-Inkugo | |
| 4C | 4C_aluminum_heat_sink | N/A | N/A | Aluminum cuboid of size approx: 100mm x 100mm x 12mm |
| 4D | 4D_screw_holder | CAD | ./01_Mechanical/P-Inkugo | |
| 4E | 4E_aluminum_heat_sink | N/A | N/A | Aluminum cuboid of size approx: 35mm x 35mm x 75mm |
| 4F | 4F_fan_holder | CAD | ./01_Mechanical/P-Inkugo | |

The upper part containing the incubation chamber is assembled as follows:

1. Take a polystyrene foam box (4A) with approximate dimensions of 240 mm × 240 mm × 250 mm and a wall thickness of approximately 45 mm.

2. Cut a square in the bottom of the box with size 75 mm × 75 mm as well as 2 holes, one for a $CO_2$ tube and one for all electrical wires.

3. Insert M3 thread inserts (#4.7) into the sensing board holder (4B) and glue it into the foam box using epoxy glue.

4. Use the aluminum block (4C) and adhere the heat sink (#4.1) using thermal paste.
5. Insert M4 thread inserts (#4.8) into the 4 (4D) 3D printed connection pieces
6. Cut 4 holes into the bottom of the foam box so that the connection pieces fit into them and glue them in using epoxy glue. Make sure they are aligned with the screw holes of the bottom layer of inkugo.
7. Next, use spray foam (#4.2) to insulate the box properly:
    a. Put some tape or foil on top of the bottom part of inkugo to protect it from the foam.
    b. Screw the two inkugo parts shown in Fig. S2C together using 4 screws (#4.6).
    c. Place the aluminum cuboid (4E) in the whole of the foam box
    d. spray the spray foam around the cuboid (do not use too much as it starts to foam up).
    e. Wait for the foam to harden.
    f. Disesamble the two halves, remove the tape/foil and trim excess foam from the polystyrene box.
8. Screw the sensing board into its holder (4B) and connect it to the electronics in the bottom layer of inkugo.
9. 3D print the fan holder (4F) and put 4 M4 nuts (#4.5) into the print.
10. Using 4 M4x16 mm screws (#4.4), screw to the fan (#4.3) to the print.
11. Place the 3D print with the fan on top of the heat sink (#4.1) and connect the fan to the electronics layer as for the inkugo presented in the main manuscript.
12. Screw the inkugo halfs together using 4 M4x10 mm screws (#4.6).
13. Finally, the cable plug (2I) can be used to plug the holes through which the cables are routed (Fig. S2D).

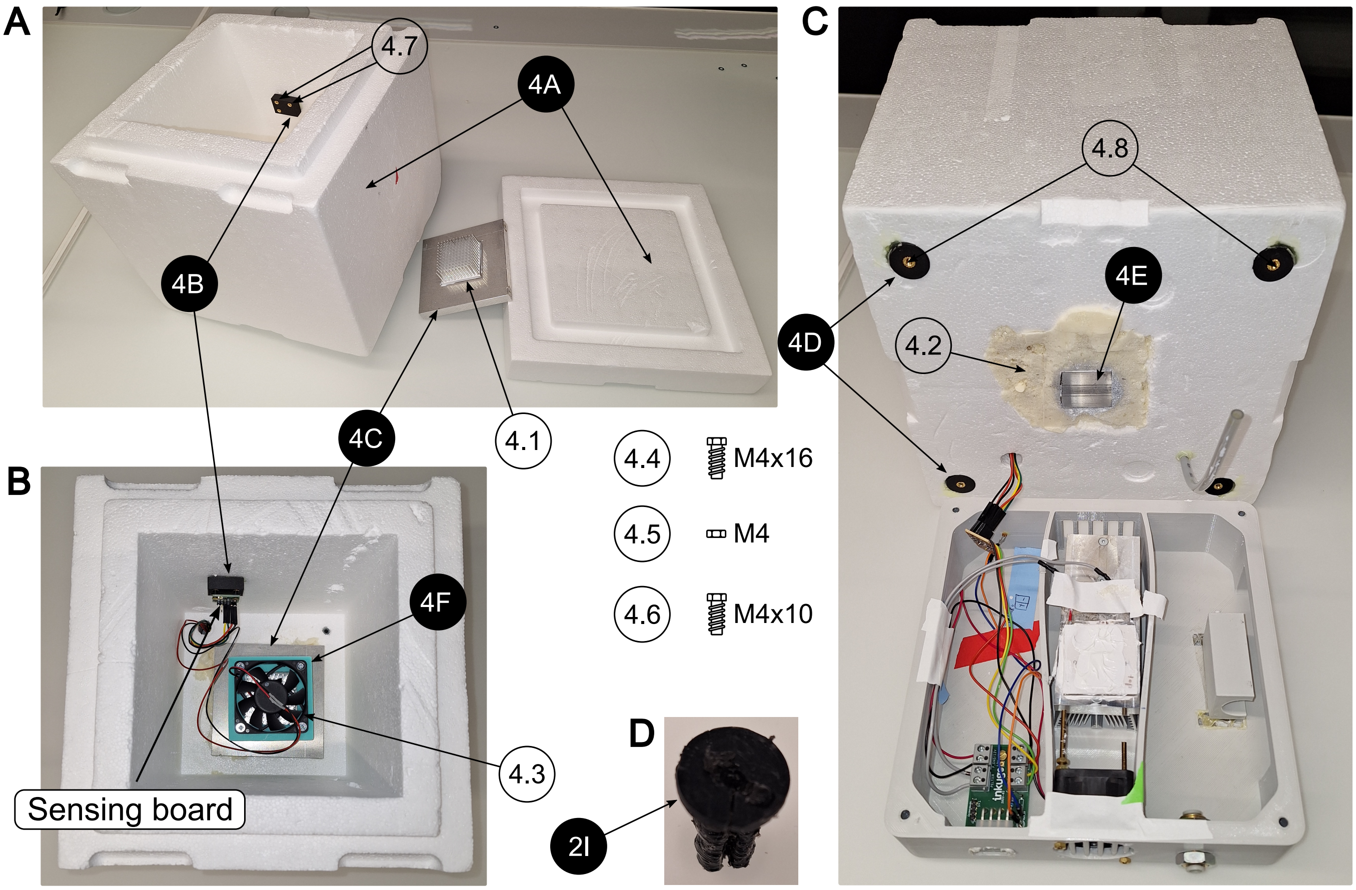


Figure S2: **Assembly of the polystyrene inkugo (p-inkugo) incubation chamber.** Design files are indicated with a black pin, other components are shown with a white pin. All components are listed in the bill of materials in the supplementary information. **A** Side view of the p-inkugo chamber without cover and parts of the cooling element. **B** Top view of the interior of the incubation chamber. **C** Side view of the disassembled p-inkugo chamber. **D** the cable plug that can be used to plug the hole used for routing the cables between the two halves of p-inkugo.

#### 2.2 Characterization

The p-inkugo can be used like the normal inkugo. To show its cooling capabilities, the system was run for approximately 10 h in cooling mode. The performance of p-inkugo is shown in Fig. S3. Starting at roughly 21 °C, p-inkugo can cool down to approximately 5 °C and stay there for a prolonged amount of time.

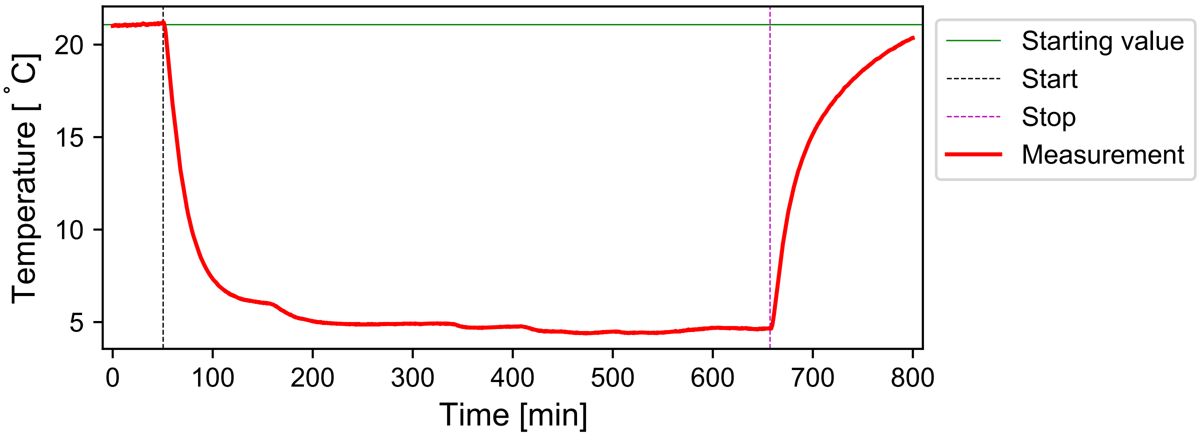


Figure S3: **Temperature curve of the inkugo system in cooling mode.** Combining the control unit with a highly insulating incubation chamber (styrofoam box) allows to establish a system suitable for cooling.
